# Post-mortem AT-8 reactive tau species correlate with non-plaque Aβ levels in the frontal cortex of non-AD and AD brains

**DOI:** 10.1101/2023.09.27.559720

**Authors:** Nauman Malik, Mohi-Uddin Miah, Alessandro Galgani, Kirsty McAleese, Lauren Walker, Fiona E. LeBeau, Johannes Attems, Tiago F. Outeiro, Alan Thomas, David J. Koss

**Affiliations:** Translational and Clinical Research Institute, Faculty of Medical Sciences, Newcastle University, UK; Biosciences, Faculty of Medical Sciences, Newcastle University, Newcastle-upon-Tyne, UK; Department of Experimental Neurodegeneration, Center for Biostructural Imaging of Neurodegeneration, University Medical Center Goettingen, Göttingen, Germany; Max Planck Institute for Multidisciplinary Sciences, Göttingen, Germany; Division of Cellular and Systems Medicine, School of Medicine, University of Dundee, Dundee

## Abstract

The amyloid cascade hypothesis states that Aβ and its aggregates induce pathological changes in tau, leading to formation of neurofibrillary tangles (NFTs) and cell death. A caveat with this hypothesis is the temporo-spatial divide between plaques and NFTs. This has been addressed by the inclusion of soluble species of Aβ and tau in the revised amyloid cascade hypothesis, however, the demonstration of a correlative relationship between Aβ and tau burden in post-mortem human tissue has remained elusive. Employing frozen and fixed frontal cortex grey and associated white matter tissue from non-AD controls (Con; n=39) and Alzheimer’s diseases (AD) cases (n=21), biochemical and immunohistochemical measures of Aβ and AT-8 phosphorylated tau were assessed. Native-state dot-blot from crude tissue lysates demonstrated robust correlations between intraregional Aβ and AT-8 tau, such increases in Aβ immunoreactivity conferred increases in AT-8 immunoreactivity, both when considered across the entire cohort as well as separately in Con and AD cases. In contrast, no such association between Aβ plaques and AT-8 were reported when using immunohistochemical measurements. However, when using the non-amyloid precursor protein cross reactive MOAB-2, antibody to measure intracellular Aβ within a subset of cases, a similar correlative relationship with AT-8 tau as that observed in biochemical analysis was observed. Collectively our data suggests that accumulating intracellular Aβ may influence AT-8 pathology. Despite the markedly lower levels of phospho-tau in non-AD controls correlative relationships between AT-8 phospho-tau and Aβ as measured in both biochemical and immunohistochemical assays were more robust in non-AD controls, suggesting a physiological association of Aβ production and tau phosphorylation, at least within the frontal cortex. Such interactions between regional Aβ load and phospho-tau load may become modified with disease potentially, as a consequence of interregional tau seed propagation, and thus may diminish the linear relationship observed between Aβ and phospho-tau in non-AD controls. This study provides evidence supportive of the revised amyloid cascade hypothesis, and demonstrates an associative relationship between AT-8 tau pathology and intracellular Aβ but not extracellular Aβ plaques.

## Introduction

The original amyloid cascade hypothesis stated that the extracellular deposition of insoluble beta-amyloid (Aβ) plaques drives intracellular tau phosphorylation, the formation of neurofibrillary tangles (NFTs), and the subsequent neurodegeneration which underlies the pathology of Alzheimer’s disease (AD) [1]. Owing to the lack of correlation between plaque burden and cognitive impairment, as well as a growing understanding of the toxicity of fibrillar and pre-fibrillar intermediate species, the hypothesis has been revised to include roles for Aβ oligomers and tau oligomers[2]. Whilst the recent outcomes of plaque clearing and Aβ oligomer targeted immunotherapies[3–5] support this revised amyloid cascade hypothesis, several inconsistencies relating to the interaction of Aβ and tau remain.

Foremost, is the spatio-temporal disconnect between the emergence and progression of Aβ plaque and tau NFT pathology. Based on the post-mortem neuropathological Thal phases of Aβ deposition and positron emission tomography (PET) imaging studies Aβ plaques originate within the neocortex, specifically within the orbito-frontal and medial parietal cortices, before spreading to the hippocampus, the brain stem and cerebellum[6, 7]. In contrast, as reflected by Braak NFT staging, tau pathology initially occurs in the entorhinal cortex and hippocampus and subsequently spreads to the lateral temporal and parietal cortices and finally to the frontal and occipital cortices[8, 9].

Cross sectional population studies further highlight the independent nature of the two hallmark pathologies, reporting that tau pathology consistent with Braak Stages I-II, occurs more readily with age than that of plaque depositions[10]. Consequently, Aβ deposition is not a prerequisite for NFT formation in aging, nor in the cases of primary tauopathies[11]. The independence of tau pathology from Aβ plaques is particularly evident in cases of primary age-related tauopathies (PART), which present with AD related NFTs in a spatial pattern consistent with Braak NFT stages up to stage IV without Aβ plaque deposition[12]. Moreover, the demonstration of prion-like spreading via tau seed templating and pathology propagation, provides a mechanistic process by which the presence of tau pathology may occur independent from the influence of Aβ plaques[13]. Such tau seed propagation of pathology likely contributes to the progression of tau pathology in many tauopathies, including AD.

Taken together, the direct causation of NFTs, purely as a consequence of Aβ plaque burden is difficult to ratify, with the differential emergence in time and space of the neuropathological hallmarks, as well as the independent occurrence of tau aggregations in other neurodegenerative conditions.

However, it remains likely that plaque deposition or rather the process of amyloid plaque formation, influences the generation of tau pathology. This is perhaps most strongly supported by numerous biochemical studies of human brain tissue, which report robust correlations with pathological Aβ and tau species[14–17]. Despite the close relationship between tau and Aβ levels in various biochemical assays, histochemical approaches frequently fail to detect such correlations. The disconnect between biochemical and histochemical analysis clearly highlights differences in the pathological species measured within the different methodological approaches.

In line with the revised amyloid cascade hypothesis[2], a range of experimental models demonstrate that soluble pathological species of both tau and Aβ exert toxic influence within the brain, evident in injection models [18–21], familial AD (FAD) [22–24], tauopathy[25–27] mouse models and cell culture approaches[28, 29]. Whilst APP centric FAD mouse models, do not develop NFTs, there is clear evidence of increased tau phosphorylation within these models. Moreover in multi-genic mice, in which a human mutant tau gene is included, APP centric mutations can accelerate and enhance NFT pathology[30]. Nevertheless, evidence for the induction of tau pathology following Aβ intracerebral delivery is sparse in non-transgenic animals[31, 32] and even in tau transgenic mice[33].

Whilst there are several possible explanations for the failure of exogenous Aβ to drive tau pathology in-vivo, one possible contributing factor is that the sole delivery of Aβ to the extracellular space may not be sufficient to drive tau pathology. Indeed, a growing body of evidence suggests that intracellular Aβ accumulation may also play a role in the pathobiology of AD[34], influencing cellular dysfunction[35–37] and tau phosphorylation[22].

Despite the potential for non-plaque Aβ to contribute to the production of tau pathology, few studies have sought to examine both biochemical and immunohistochemical quantification of Aβ and tau pathology within the same cases, thus, allowing for a direct comparison between the correlative strength of total Aβ, plaque Aβ and intracellular Aβ with tau pathology. This current study aimed to quantify such parameters across cases to establish potential correlative relationships between pathological species in order to further understand the influence that Aβ, in various forms, may have upon the regional generation of tau pathology, both in neuropathologically confirmed cases of AD as well as non-AD control cases.

## Methods

### Human post-mortem brain tissue

A study cohort of post-mortem human brains from clinico-pathologically classified AD (n=21) and non-neurodegenerative control cases (Con, n=39) was obtained from the Newcastle Brain Tissue Resource (NBTR). AD Subjects had been clinically assessed during life prior to brain tissue donation and diagnosed with dementia due AD. Control cases similarly had been assessed during life and at the time of death did not have dementia. The final clinico-pathological diagnoses were established by combining clinical neuropathological data reviewed at regular meetings involving JA and AT. Neuropathological diagnoses were based on assessment of brain tissue according to the National Institute of Ageing – Alzheimer’s Association (NIA-AA) criteria[38], including Braak NFT staging[39], Thal phases [6], and Consortium to Establish a Registry for Alzheimer’s Disease (CERAD) scoring[40], as well as Braak LB stages[41], and Newcastle / McKeith Criteria [42, 43] (Table 1 and S1 for full details).

**Table 1.**
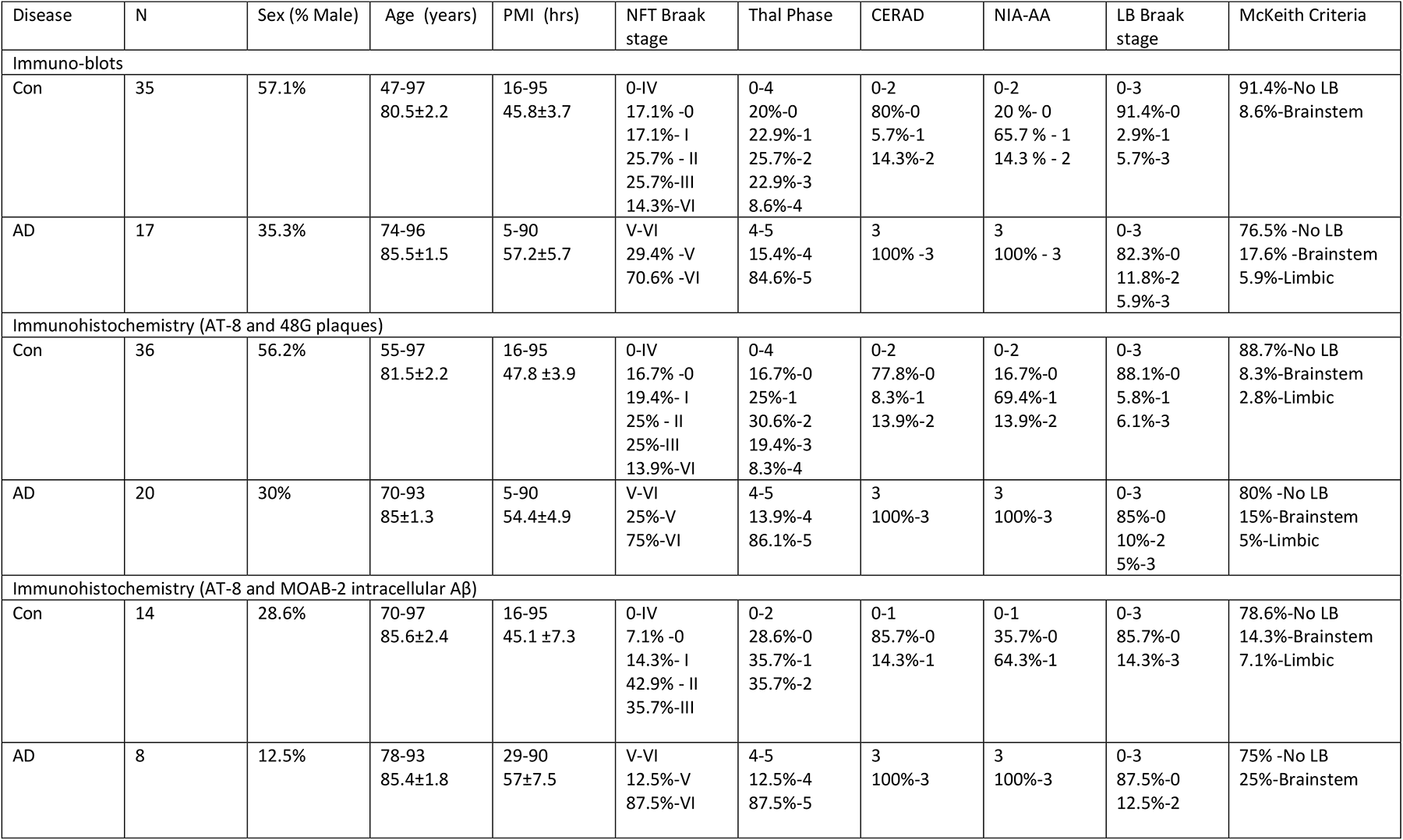
Post-mortem human tissue cases and use. Human cases use for immuno-blots and immunohistochemistry for plaques and AT-8 as well as intracellular Aβ and AT-8 are listed. Case are separated by disease classification according to non-diseased controls (Con) and Alzheimer’s disease (AD). Case numbers (n), sex, age, post-mortem interval (PMI), neurofibrillary tangle (NFT) Braak stage, Thal phase, Consortium to Establish a Registry for Alzheimer’s Disease (CERAD), the National Institute of Ageing – Alzheimer’s Association (NIA-AA) criteria, Lewy body (LB) Braak stage and McKeith criteria are provided. For age and PMI both range and mean ±SEM are provided. For numerical scores of pathology, range and percentage composition are given. For CERAD scores, negative (neg), A and B reported. For NIA-AA, not, low and intermediate (inter) risk for Alzheimer’s disease. For McKeith criteria, only percentage composition is given, where cases free of LBs (No LB), brainstem, Limbic and neocortical (Neo) predominate are indicated.

For histology and tissue micro-array (TMA; see below), tissue sections were prepared from the right hemisphere of the brain and fixed for 4-6 weeks in 4% paraformaldehyde. Corresponding frozen frontal grey (GM) and white matter (WM) tissue (Brodmann’s area (BA) 9) was obtained from the left hemisphere, dissected in a coronal plane and snap frozen between copper plates at -120°C prior to being stored at -80°C. Due to limitations in tissue availability, it was not possible to obtain both fixed and frozen tissue for all cases (see Table 1 and S1 for full details). Comparative analysis of age and post-mortem interval (PMI) between disease groups determined no significant difference in either measure (p>0.05).

### Tissue lysis

∼250mg of frozen frontal tissue was electronically homogenised 1:10 (W/V) in 0.2M tetraethyl ammonium bicarbonate (TEAB, pH 7.2, Sigma) with 1% SDS, containing protease (1 per 10ml, Complete, Roche) and phosphatase inhibitors (1 per 10mls, PhosSTOP, sigma) using an Ultra-turrax T10 homogeniser (5 mm diameter probe; 30,000rpm) for 15 sec. Lysate were aliquoted and stored at -80°C, prior to use.

#### Immunoblot quantification of AD markers

Dot blots were conducted for total Aβ and AT-8 phospho-tau in both grey and white matter samples. The protein concentration of GM and WM crude lysate were adjusted to 0.5µg/µl as per Bradford assay and dotted directly to a nitrocellulose membrane at 10μl (5µg/dot) and left to dry for 20 minutes before further processing. The membranes, briefly washed in Tris-buffered saline (TBS; in mM; 50 Trizma base, 150 NaCl, pH= 7.6) prior to being blocked in 5% milk powder containing Tris-Buffered Saline with 0.1% Tween 20 (TBST) at room temperature for 1 hour. After blocking, blots were rinsed in TBS washing buffer 3 times for 5 minutes each. Membranes where subsequently placed in primary antibody solution (TBST, 5% bovine serum albumin and 0.05% sodium-Azide) containing either MOAB-2 (1:1000, Cat# M-1586-100, Biosensis) for the detection of Aβ or AT-8 (1:1000, Cat# AB_223647, Thermofisher) for phospho-tau and incubated overnight at 4°C. The membranes were then washed in TBST before being incubated for 1 hour at room temperature in horse radish peroxidase conjugated goat anti-mouse secondary IgG antibody (TBST+5% milk powder+1:5000 dilution) prior to repeated washing before being development. Immunoreactivity was visualized via enhanced chemiluminescence (1.25 mM Luminol, 25μl of 3%H202 and 50μl coumaric acid was incubated for 1 minute). The signal was captured by using digital western blot camera, with high sensitivity and auto exposure being selected. The images were saved as 8-bit for illustration and 16-bit for quantification. Total protein loading was determined via Ponceau S general protein stain (0.1% Ponceau S (w/v) and 5.0% acetic acid (w/v) in ddH_2_O water) and resulting loading staining captured.

### Immuno-blot quantification

Immunoreactivity of enhanced chemiluminescent (ECL) luminol blots and Ponceau S-stained blots was quantified from 16-bit digitized images based on area under the curve measurements as computed by ImageJ (Ver 1.53e, NIH, USA). Normalization of immunoblot intensity values were then performed using total protein adjusted values. The 52 samples of human frontal cortex GM and WM were processed in 4 separate batches and each batch normalized to the mean value of control cases (each blot containing >3 Br 0-IV control cases) prior to pooling values between blots.

### Immunohistochemical quantification of Aβ plaques and phospho-tau (AT-8)

Regional quantification of the Aβ plaque and AT-8 phospho-tau load within the frontal cortex (BA9) via TMA slides, as described previously[44]. Sections (6 µm, thick) were cut from paraffin embedded TMA blocks tissue blocks comprising of cylindrical tissue cores taken from multiple brain region specific blocks and mounted on glass slides. Slides containing a 3 mm diameter samples of BA9 frontal cortex were baked at 60°C for 1hr prior to being dewaxed in xylene, rehydrated in descending concentration of ethanol (5 mins immersion) and washed in TBS. Slides intended for phospho-tau staining were treated with microwave assisted antigen retrieval (800 W, 10 mins) in citrate buffer (10 mM Citric acid, 0.05 % Tween 20, pH 6) and those intended for Aβ plaque staining were submerged in 90% Formic acid for 1hr at RT, before endogenous peroxidases were quenched in H_2_O_2_ (3%, 20 mins submersion). Following consecutive washes in TBS and TBST, slides were incubated with either mouse 4G8 (1:16000, Cat# SIG-39200, Covance) or anti-AT-8 (1:4000) in TBS for 1 hr and immunoreactivity visualised via the MENAPATH HRP polymer detection kit (Menarini diagnostics, Wokingham, UK) and 3,3’-Diaminobenzidine (DAB) chromogen with appropriate TBS and TBST washes performed between steps. Slides were co-stained with haematoxylin prior to being dehydrated in ethanol, cleared in xylene and mounted in dibutylphthalate polystyrene xylene (DPX).

Stained BA9 frontal cortex samples were imaged at x100 magnification with a semi-automated microscope (Nikon Eclipse 90i microscope, DsFi1 camera and NIS elements software V 3.0, Nikon). For each case, multiple images were captured to form a 3x3 image grid with 15% overlap in adjacent images, such that an area of 1.7mm was sampled from each case.

Following visual quality control inspection and the application of regions of interest (ROI) to exclude areas of tissue folds and tears, a consistent restriction threshold for 4G8 (R50-180, G20-168, and B8-139) and AT-8 (R25-170, G27-156, B11-126) was applied producing a binary signal image from which the percentage area of immunoreactivity could be acquired. For the quantification of Aβ plaques, 4G8 images were further processed by means of size exclusion, restricting object detection to >100µm^2^, thus avoiding inclusion of intracellular APP and Aβ.

### Immuno-fluorescent histochemical analysis of intracellular Aβ and phospho-Tau (AT-8)

Paraffin embedded tissue blocks of the frontal cortex BA9 were used to prepare sections (6 µm thick) for the purpose of multiplex intracellular Aβ and phospho-tau fluorescent staining. Slide mounted frontal cortex sections were baked at 60°C for 1hr, dewaxed and rehydrated and subjected to antigen retrieval in citrate buffer and formic acid treatment (as above). Slides were then blocked in TBST containing 10% normal goat serum for 1hr at RT and incubated in mouse IgG2b anti-MOAB-2 and mouse IgG1 anti-AT-8 (1:500, for both) overnight at 4°C, prior to incubation in secondary antibodies (goat-anti mouse IgG1 – Alexa 488 and goat-anti mouse IgG2b-Alexa 594, 1:1000 for both, Invitrogen). Endogenous tissue fluorescent was quenched via post-staining treatment with Sudan Black (0.01%, 70% ethanol, 5mins submersion) before slides were coverslipped with DAPI-containing Prolong Diamond Mounting media (Fisher Scientific). In a subset of slides, the limited co-localization of MOAB-2 labelled Aβ and APP was established, staining sections with mouse-IgG2b anti-MOAB-2 and Rabbit-anti-APP (1:500, Cat# ab15272, abcam) and appropriate secondary antibodies. Fluorescence antibody labelled sections were imaged via a wide-field fluorescence microscope system (Nikon Eclipse 90i microscope, DsQi1Mc camera and NIS elements software V 3.0, Nikon).

One section per case was examined at 400x magnification with 3 images per grey matter and white matter region selected at random. As these images were used for quantification of intracellular Aβ, excluding Aβ-plaques, any region selected which contained multiple plaques was excluded and another region selected. ROI were manually applied to each image and folds, tears and plaques were excluded, before images were converted to grey scale and a consistent threshold applied to generate a binary image from which percentage area of immunoreactivity was determined. The mean percentage area of immunoreactivity was calculated per grey and white matter area per case.

### Data analysis

Data were subjected to Shapiro-Wilk normality test for normal distribution, prior to statistical comparison between control and AD cases using a non-parametric Mann-Whitney U test (GraphPad Prism Ver. 5). In SPSS, two-tailed Spearman’s correlation was used for correlation analysis. Given the association of increasing Braak stage with age, all correlations with Braak staging were performed with partial correlations controlling from age. A series of one-tailed t-test were performed to identify the initial stage at which measures were significantly elevated from Braak 0 pathological controls. For all analysis, p<0.05 was considered as statistically significant, with increasing statistical reliability for p<0.01, p<0.001 and p<0.0001.

## Results

### Biochemical analysis of Aβ and phospho-tau pathology

In order to limit the potential confounding influence of age-related Aβ independent tau pathology as seen in PART[12], Brodmann’s area 9 of the superior frontal cortex, a region which does not develop NFTs until late stage of AD related disease progression (Braak NFT V-VI) was selected for investigation. Using a native state dot-blot quantification, crude tissue lysates of grey matter of this region as well as the associated white matter of AD (n=17) and non-AD controls (n=35), were probed for Aβ via the non-APP cross-reactive MOAB-2 antibody and for tau pathology using the phospho-tau specific antibody AT-8.

When considered purely based on the neuropathological diagnosis of either non-AD and AD, levels of AT-8 phospho-tau were elevated in AD cases, both in GM (13.24±3.3-fold cf. non-AD, p<0.001, fig1 a.i + b.i) and WM (9±2.6 fold cf. non-AD, p<0.001, fig1 a.i + b.i). Despite a numerically higher mean within the GM compared to the WM, there was no statistically significant difference between the magnitude of increase between GM and WM in AD cases (p>0.05). Similarly, when Aβ levels were examined based on the neuropathological cohort stratification of Non-AD and AD, elevations were apparent within the GM (3.35±0.59 fold c.f non-AD, p<0.01, fig1 a.ii + c.ii) and WM (2.56±0.56 fold c.f non-AD, p<0.05, fig1 a.ii + b.ii) of AD cases. Again, no difference between the magnitude of increase within AD cases relative to control cases was observed between GM and WM (p>0.05).

Such an outcome from the analysis of phospho-tau and Aβ between AD and non-AD cases is not surprising but serves to validate the use of dot-blots to measure biochemical changes in phospho-tau and Aβ.

To further place the observed changes of tau and Aβ within the context of disease progression, the cohort was subdivided into their respective Braak stages (0-VI) and the association of crude lysate measures of AT-8 phospho-tau and Aβ with disease progression was determined (Fig 1 c.i + ii). Phospho-tau AT-8 immunoreactivity correlated with increasing Braak stages when considered across the entire cohort within the GM (r=0.7, p<0.001) as well as in the WM (r=0.67, p<0.001). Additionally, a modest but significant correlation between Braak NFT stage and age was reported (r=0.34, p<0.05) although age did not correlate with biochemical measures of AT-8 (p>0.05, data not shown). Correlations between AT-8 immunoreactivity and Braak NFT remained when adjusting for age (r=0.62 and r=0.57 for GM and WM respectively, p<0.001 for both, fig 1 c). Interestingly when probed for the stage at which phospho-tau levels were significantly elevated from that of “pathologically-free” Braak stage 0 cases, Braak stage IV was indicated (p<0.05, for both GM and WM). Braak stage IV being the stage prior to gross affection of the frontal cortex with NFTs. Furthermore, when split according to neuropathological diagnosis, and controlled for age, a significant correlation was observed in the non-AD control group in GM (Fig 1 c.i, r=0.47, p<0.01) and WM (Fig 1c.i, r=0.43, p<0.05), but not in AD cases (Fig 1 c.i, p>0.05). Together the data suggest that in non-AD control cases, AT-8 phospho-tau increases within the frontal cortex in line with the intra-regional spatial AT-8 positive NFT progression, somewhat independent of age. Thus, initial levels of frontal cortex AT-8 positive phospho-tau may be regionally generated.

**Figure 1.**
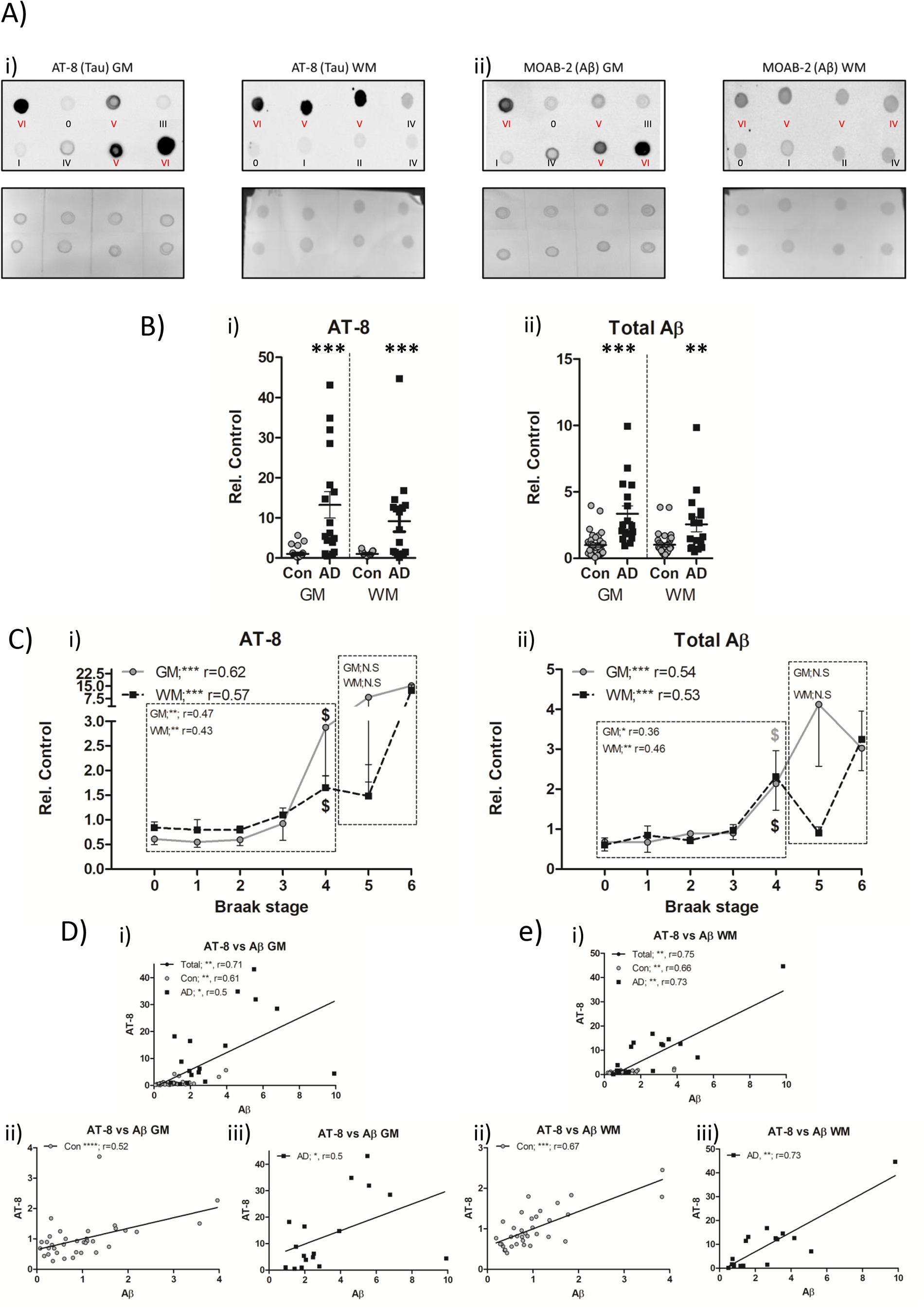
Biochemical quantification of AT-8 phospho-tau and Aβ in the frontal cortex of non-AD and AD cases. A). Example dot blots of AT-8 (i) and MOAB-2 (Aβ; ii) immunoreactivity and associated ponceau total protein stain, produced from crude tissue lysates of frontal grey (GM) and white matter (WM) in control (Con; black lettering) and Alzheimer’s disease (AD; red lettering) cases. Braak NFT stage of each sample is shown. B). Comparison of mean AT-8 (i) and Aβ (ii) immunoreactivity between control (Con) and Alzheimer’s disease (AD) cases in the GM and WM C).Association of AT-8 (i) and Aβ (ii) immunoreactivity with Braak NFT stages across the cohort in GM and WM. Correlative analysis (Spearman’s r) is shown for when analysis as a single group or when separated into Con and AD groups. Combined (i), Con(ii) and AD (iii), linear correlations between AT-8 and Aβ in the GM (D) and WM(E). Immunoreactive shown as relative to control (Rel. Control). *=p<0.05, **=p<0.01, ***=p<0.001 and ****=p<0.0001. $ denotes initial Braak NFT stage at which immunoreactivity is significantly elevated from Braak 0 controls.

Following a similar line of investigation for the accumulation of Aβ in relation to disease progression, Aβ levels were correlated with individual Braak NFT stages. Across the entire cohort, robust correlations were reported for both the GM (r=0.67, p<0.001, fig 1 d) and WM (r=0.6, p<0.001, fig 1 d). Again, correlations remained when controlling for age (r=0.54 and r=0.53 for GM and WM respectively, p<0.001 for both). In line with observations of AT-8 phospho-tau, individual comparisons with Braak 0 cases reported an initial significant elevation from the “pathologically-free” baseline at Braak 4, in GM and WM samples (p<0.05). Consistent with similar measures of AT-8 phospho-tau, when divided into non-AD and AD categories, increasing total Aβ correlated with progressive Braak stages in the non-AD group in both the GM and WM (r= 0.36, p<0.05 and r=0.46, p<0.01 in GM and WM respectively, fig 1 d) and not in the AD group (p>0.05). Similar to correlation with Braak staging, all other neuropathological classifications also reported robust correlations when considered as a single cohort (See S.table 2). When considering Non-AD controls only, significant correlation with AT8 phospho-tau was only apparent with CERAD scores for GM and WM (r=0.4, p<0.05 and r=0.47, p<0.001, respectively) whilst GM and WM scores for Aβ correlated with Thal, CERAD and NIA-AA (r=0.38-0.52, p<0.05, see S.table 2 for full details). In terms of AD cases, correlations were apparent for all biochemical measures with all neuropathological scheme (see S.table 2).

Collectively, the data suggest a robust relationship of AT-8 and Aβ with disease progression within the GM and WM of the frontal cortex, occurring not only in confirmed cases of AD but also non-AD controls. In the frontal cortex, intra-regional pathology progresses in accordance with global AD-related brain tau pathology as quantified by Braak NFT stages and this was most evident in non-AD controls.

Independent of global brain AD related pathological changes, a critical element to regional pathology is to determine whether biochemical measures of tau pathology and Aβ correlate on a case-by-case basis, as the disconnect of histochemical tau and Aβ pathological hallmarks has long been a major caveat to the amyloid cascade hypothesis. Indeed, a robust correlation between biochemical measures of AT-8 phospho-tau and total Aβ measures was observed when considered as a single cohort (Non-AD + AD cases) in the GM (r=0.75, p<0.001, fig 1 d.i) and in the WM (r=0.67, p<0.001, fig1 e.i). Remarkably, when examined separately within Non-AD controls and AD cases, correlations between AT-8 phospho-tau and Aβ were apparent in both control cases (GM; r=0.52, p<0.001 and WM; r=0.67, p<0.001, fig 1 d ii + e ii) as well as AD cases (r=0.5, p<0.05 and r=0.73, p<0.01 in GM and WM, fig 1 d iii + e iii).

### Histochemical quantifications of AT-8 phospho-tau and Aβ plaques

In order to establish if the biochemical derived relationship of increased in Aβ immunoreactivity conferring increased in AT-8 immunoreactivity was primarily driven by an association of Aβ plaques with AT-8 phospho-tau, semi-quantitative immunohistochemistry analysis was performed (Fig 2 a, and see S.table 1 for details). Based on % area coverage, AT-8 immunoreactivity, a composite of NFTs and NTs reported a marked ∼ 100 fold increase in % coverage in the frontal grey matter of AD cases compared to controls (7.6±2.5% vs. 0.07±0.02%, in AD and non-AD cases respectively, p<0.001, fig 2 b.i). Equally, quantification of the % area coverage of Aβ plaques between AD cases and non-AD controls, also, unsurprisingly reported a significant increase with the AD cases (14.26±1.7% cf. 3.55±1%, p<0.001, fig 2 b.ii).

**Figure 2.**
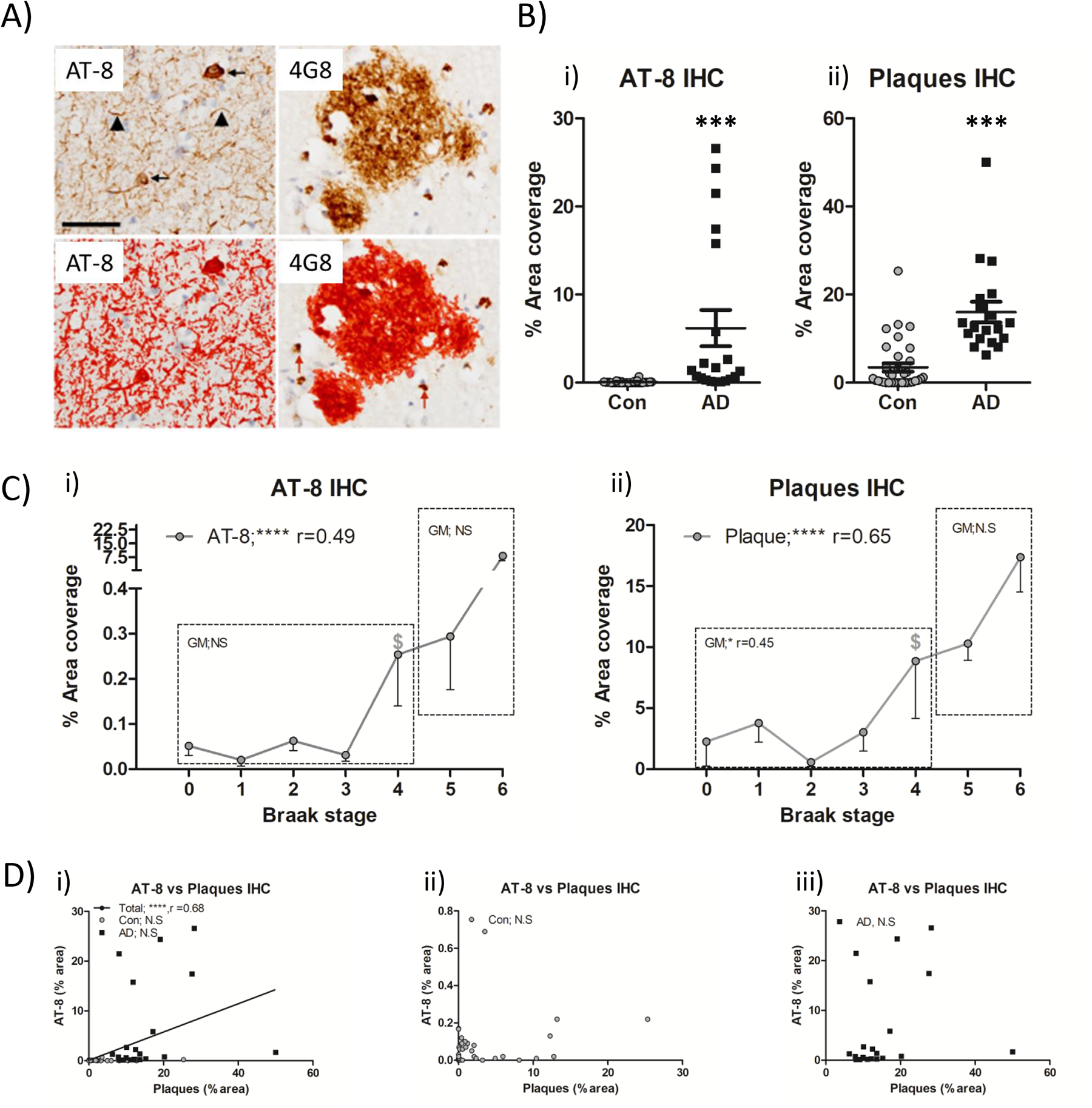
Immunohistochemical quantification of AT-8 tau and Aβ plaque burden in the frontal cortex of non-AD and AD cases. A). Example micrographs of DAB based AT-8 and 4G8 (Aβ) immunoreactivity, area of quantification following threshold application shown in red. Note the size exclusion in this parameter of intracellular 4G8 labelling to negate potential APP cross-reactivity. B). Quantification of % area coverage of AT-8 (i) and plaque (ii) immunoreactivity in control (Con) and Alzheimer’s disease (AD) cases. C). Association of % area coverage of AT-8 (i) and plaques (ii) with Braak NFT stage with correlative analysis (Spearman’s r) shown. Combined(i), Con(ii) and AD(iii), linear correlations between AT-8 and plaques in the GM (D). N.S = not significant, ***=p<0.001 and ****=p<0.0001. $ denotes initial Braak NFT stage at which immunoreactivity is significantly elevated from Braak 0 controls. Scale in a = 20 µm.

When measures were considered in relation to progressive Braak NFT staging, AT-8 phospho-tau (r=0.55, p<0.001, fig 2 c.i) and Aβ plaques (r=0.67, p<0.001, fig 2 c.ii) strongly correlated with Braak stage, following a correction for age. Again, a significant elevation in AT-8 phospho-tau and Aβ plaque coverage from “pathologically-free” Braak stage 0 cases was reported at Braak Stage IV (p<0.05), in line with observations from biochemical measurements. When controlling for age, no significant correlations were observed for AT-8 phospho-tau or Aβ plaques for either non-AD or AD groups (p>0.05). Further analysis with additional neuropathological classification reported strong correlations with Thal and CERAD and NIA-AA when considered as a single cohort and in control only (S.table 3). Collectively, the data largely suggests that correlative relationships reported within the overall cohort likely stems from group effects driven by the general increase of pathological hallmarks between non-AD and AD groups and not as incremental regional increase in line with progressive Braak stage. Such observations contrast with the findings of the biochemical investigation.

Equally correlations between IHC quantified Aβ and AT-8 phospho-tau also reported a correlation only when the data were analysed as a single cohort combining non-AD controls and AD cases (r=0.65, p<0.001, fig 2 d) and not when examined as a separate data set of non-AD controls or AD cases (p>0.05, for both, fig 2 dii+iii). Nevertheless, biochemical measures correlated with respective histochemical measures for AT-8 and Aβ (see Stable 4). Thus, despite the contradiction between biochemically measured association of AT-8 phospho-tau and total Aβ and histologically measured AT-8 phospho-tau and Aβ plaque load, the measures by differing methodological means are somewhat inter-related.

### Quantification of intracellular Aβ

The absence of a correlation between IHC measures AT-8 and plaques (fig 2), despite a robust correlation with of biochemical AT-8 and total Aβ measures from crude tissue lysates (fig 1), suggests the possible inclusion of additional Aβ sources within the biochemical quantification. Accordingly, the application of MOAB-2 Aβ antibody to fixed post-mortem human brain tissue sections, labelled both extracellular plaques and intracellular pools of Aβ (fig 3). Comparisons of APP and MOAB-2 Aβ labelling, demonstrated a clear distinction in subcellular and plaque labelling in both GM (fig 3 a) and WM (fig 3 b). Whilst there is a degree of overlap as would be expected given the spatial limitation and the production of Aβ from APP, co-localisation is absence in the majority of puncta (fig 3; white arrows), confirming the specificity of MOAB-2 for the labelling of Aβ, as previously reported[45, 46]. Across a subset of the cohort, both in the GM and WM, a progressive increase in intracellular Aβ staining in line with increase AT-8 immunoreactivity was observed (fig 4.a). When intracellular Aβ was quantified according to % area coverage, a significant increase in the levels was observed in AD cases compared to controls (fig 4b.i, p<0.001 in GM and p<0.05 in WM). Similarly, AT-8, as measured by immunofluorescence was again elevated in AD cases (fig 4 b ii, p<0.001 in GM and WM). Both AT-8 (r=0.62, p<0.01 in GM and r=0.5, p<0.05 in WM) as well as intracellular Aβ (r=0.61, p<0.01 in GM) correlated with Braak stage when controlling for age. Equally, strong correlations were observed with Thal, CERAD and NIA-AA also (see S.table 5). No correlations with any neuropathological classifications were reported when the cohort was spilt according to Non-AD and AD groups. Nevertheless, correlative measures between intracellular Aβ and AT-8 phospho-tau reported a significant relationship across all cases (in GM, r=0.76 and in WM, r=0.71, p<0.01 for both, fig 4 c i + d i,) and in control cases only (in GM, r=0.82 and in WM, r=0.76, p<0.01 for both, fig 4 c ii + d ii). Interestingly when considering only the AD cases, a significant inverse correlation of intracellular Aβ with AT-8 phospho-tau in the GM, was apparent (r=-0.74, p<0.05, fig 4 c iii), although no correlation was reported in the WM (fig 4 d iii). As with IHC quantifications of plaques and AT-8 as per TMA slides, measures of intracellular Aβ and AT-8 here, also correlated with biochemical measures when considered as a single group, although no significant correlations were observed when split into Control and AD categories (S. table4).

**Figure 3.**
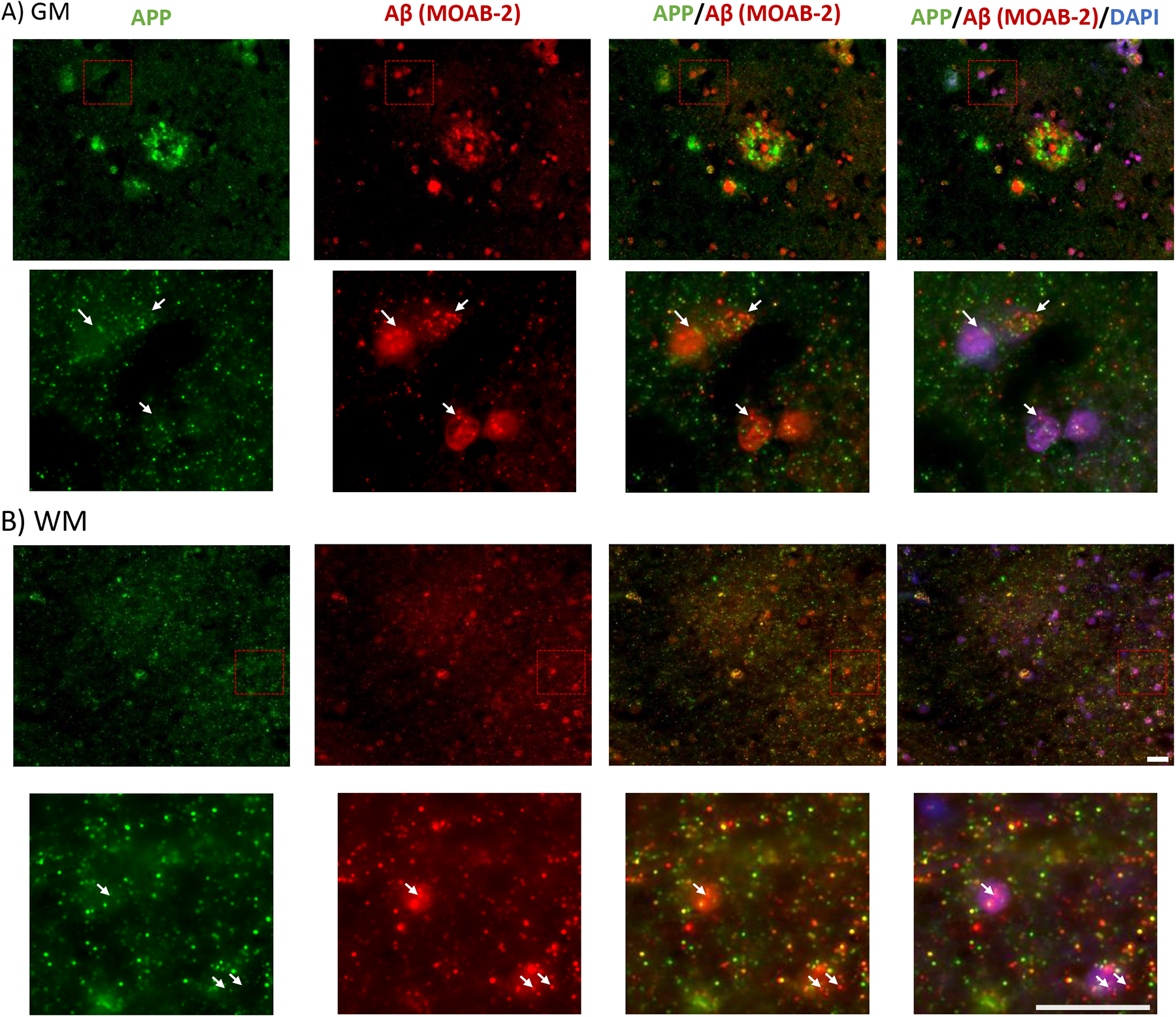
Immunohistochemical distinction between MOAB-2 labelled Aβ and APP immunoreactivity in the frontal cortex of an AD case. Example micrographs of APP (N-terminal-APP antibody) and Aβ (MOAB-2) from an AD case, in the grey (GM; a) and white matter (WM; b). Note the distinctive labelling of subcellular pools within insert (white arrows) and differential labelling of plaques (in a) Scale =20 µm.

**Figure 4.**
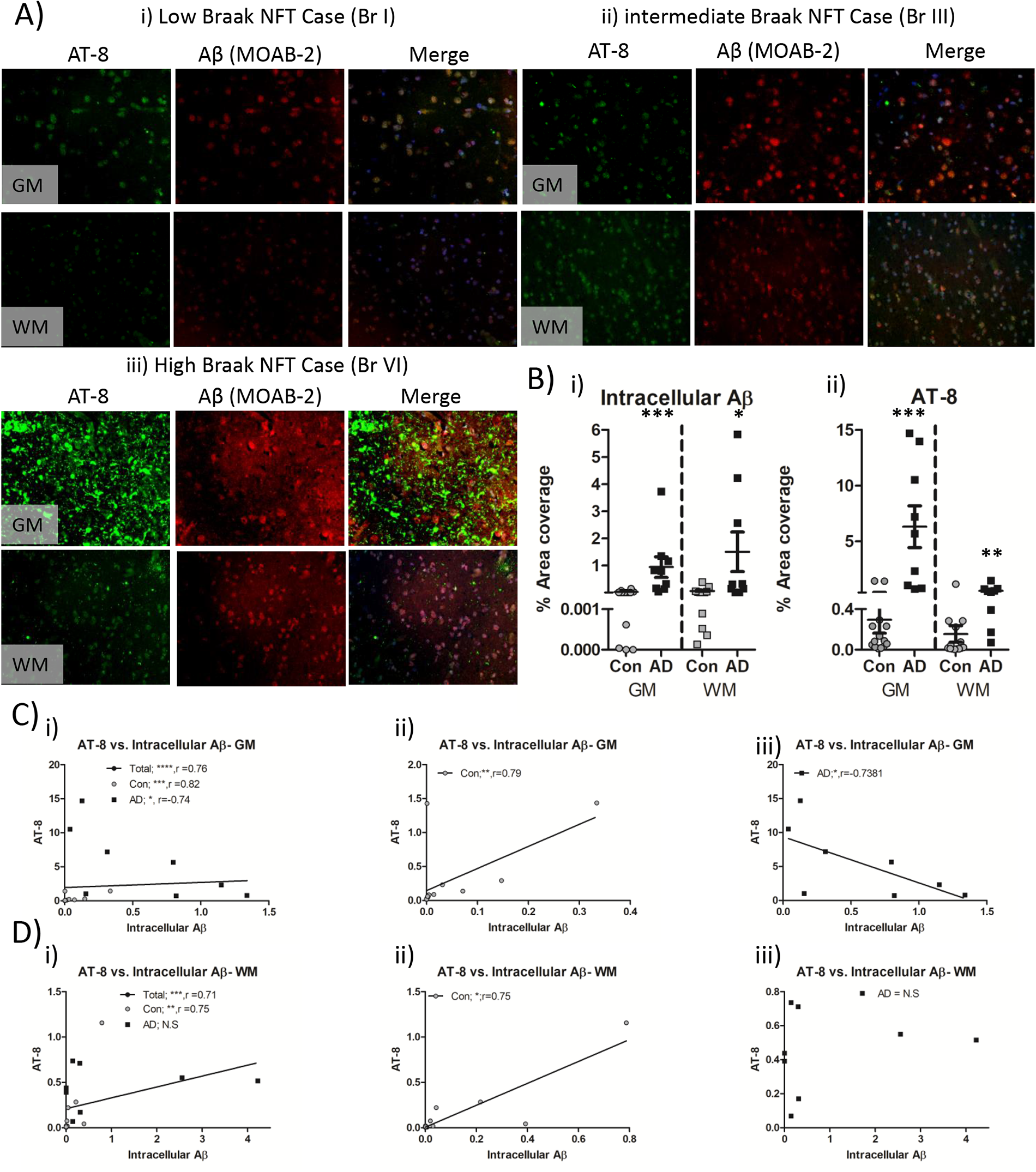
Immunohistochemical quantification of intracellular Aβ and AT-8 tau in the frontal cortex of non-AD and AD cases. A). Example micrographs of AT-8 phosphorylated tau and MOAB-2 labelled intracellular Aβ in GM and WM of low (i), intermediate (ii), and high (iii) Braak stage cases. B). Quantification of intracellular Aβ (i) and AT-8 phospho-tau (ii) expressed as percentage area coverage in control (Con) and Alzheimer’s disease (AD) cases. Combined(i), Con(ii) and AD(iii), spearman’s correlations (r) between AT-8 and plaques in the GM (C) and WM (D). N.S = not significant, *=p<0.05, **=p<0.01, ***=p<0.001 and ****=p<0.0001. Scale =20 µm.

Collectively quantification of the intracellular Aβ and its correlation with AT-8 phospho-tau appears to support the biochemical findings of a close relationship between AT-8 and Aβ, particularly in non-AD control cases. However is should be noted that spatial colocalization of AT-8 and intracellular Aβ was not common and appeared to be the exception rather than the rule (fig 5), instead here we find a correlation of regional AT-8 burden with regional intracellular Aβ.

**Figure 5.**
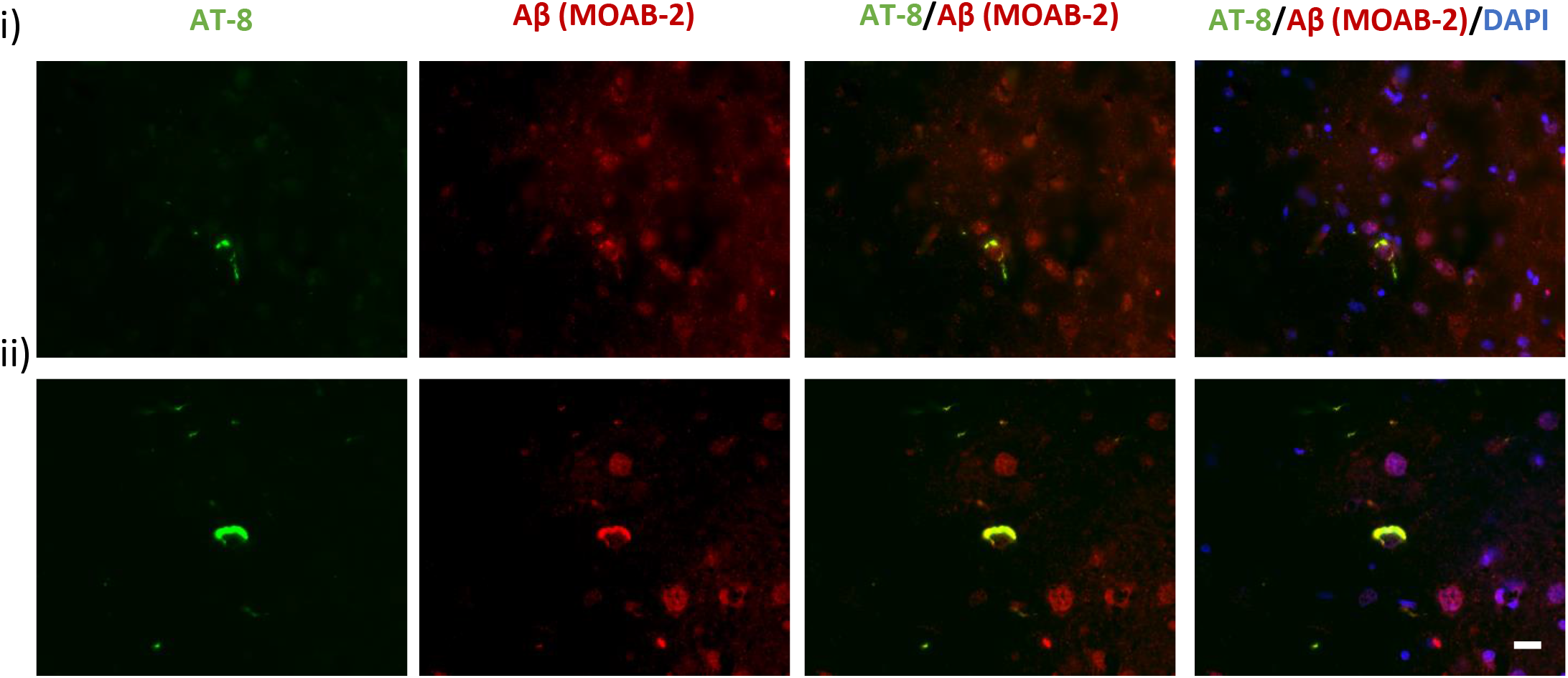
Rare instances of intracellular Aβ and tau colocalization. Example micrographs demonstrating an overlap of AT-8 and Aβ immunoreactivity within cells from a non-AD control Braak stage IV. Scale=10 µm.

## Discussion

Collectively this study reports an apparent correlation between biochemical Aβ and AT-8 phospho-tau measures. Such a relationship was not reproduced when comparing IHC based quantification of Aβ plaques and AT-8 phospho-tau but was observed when considering IHC measures of phospho-tau and intracellular Aβ. These correlative relationships are perhaps surprisingly strongest within the non-AD control group and are evident in both grey and white matter. Together the data suggests a close relationship between non-plaque Aβ and tau, which is at least partially due to the accumulation of intracellular Aβ and its potential influence on tau phosphorylation.

### Regional correlation of tau and Aβ

A long-standing critique of the amyloid cascade hypothesis has been the disconnect between NFT and Aβ-plaque burden within a given brain region of either non-AD controls or indeed AD cases [47, 48]. However many biochemical approaches have previously found correlations between Aβ, either total Aβ or specifically Aβ1-42, with a range of phosphorylated and oligomeric tau markers within the GM of a given cortical region[14–17]. Furthermore, disease dependent changes in white matter Aβ levels have also been previously observed using Aβ_40_ and Aβ_42_ ELISAs[49] and like-wise hyperphosphorylated tau has also been observed biochemically within the white matter of AD cases[13, 50]. Here we observed that levels of AT-8 reactive phospho-tau and Aβ increased in non-AD controls in line with Braak stage progression, in both the frontal grey and white matter and exhibited a positive correlation between Aβ levels and AT-8 when measured biochemically. Whilst our observations cannot determine causality, our findings are consistent with many *in-vitro* experiments in which the application of Aβ to various cellular preparations results in downstream tau phosphorylation[28, 51, 52]. Moreover, support for the interaction of Aβ with tau pathology, can be gained from studies reporting that interventions targeting Aβ levels consequently reduce tau pathology both *in-vitro* and *in-vivo* models[53, 54] as well as in biofluids obtained from human clinical trials[3, 4, 55].

When considering the association of Aβ and tau phosphorylation within non-AD controls, our findings are similar to the observation of a previously seen correlation between Aβ_1-40_ level and p-181 tau in the CSF of control cases[56]. However here we can extend this finding to show that within the frontal cortex there is a regional dependence between Aβ levels and tau pathology. As work by others have reported no or minimal seed competent tau within the frontal cortex in non-AD controls [50], the source of pathology within the frontal cortex is likely to mainly be intraregional, with minimal influence of extra-regional spread. Such intraregional dependence would be consistent with the strong correlation between Aβ and AT-8 signals within the non-AD controls, which is somewhat weakened in AD cases, when seed competent tau species have presumably invaded the frontal cortex[13], the self-propagation of tau pathology thus diminishing the relative contribution of Aβ to the production of phospho-tau species. Interestingly, a linear relationship between regional soluble tau phospho-species and Aβ_1-42_ has previously been reported alongside a bimodal relationship within the insoluble fraction, implying a weakening of the relationship between Aβ_1-42_ and tau phosphorylation once aggregated [17]. Thus, given that this present study did not seek to distinguish between soluble and insoluble pathology, the increased representation of insoluble tau pathology within lysates from AD cases, likely further explains the weakening of the linear relationship between Aβ and tau phosphorylation in AD cases.

Accordingly, no linear correlation was observed between plaque load and AT-8 load when measured histologically. Such differences may relate to the presumed loss of extracellular soluble Aβ and exclusion of intracellular Aβ pools, as is common practice when assessing Aβ burden as part of a neuropathological assessment[44].

### Intracellular Aβ

Historically, intracellular Aβ has been problematic in its quantification, largely due to the cross-reactivity of Aβ antibodies with APP and other intermediate APP metabolites. Howecer, several commercial Aβ antibodies are available, including MOAB-2 which shows no cross-reactivity with APP under many conditions [45, 46].

Although often overlooked, the production of Aβ occurs intracellularly following endosomal APP cleavage via β-secretase [57] and sequential γ-secretase processing within either Golgi [58] or lysosomal [59] compartments. Whilst the majority of Aβ is trafficked to the extracellular space, age-related changes in the relative production of Aβ peptide length [60] and disease alterations to trafficking mechanisms, such as Rab GTPases [61, 62], may act synergically to enhance the retention of intracellularly produced Aβ and or indeed its reuptake, leading to its intracellular accumulation [63]. Accordingly, post-mortem examination of the entorhinal cortex and hippocampus of non-diseased non-AD cases, suggests an increase in intracellular Aβ in line with increasing age [64, 65] and furthermore AD animal models also show an age-related accumulation of intracellular Aβ [46, 66].

### Intracellular Aβ and tau

Here, when selectively measuring intracellular Aβ, a positive correlation between Aβ and AT-8 in the frontal cortex of non-AD controls was observed. This is consistent with biochemical measures of frontal cortex lysates in the same cases as present here or indeed previously reported biochemical measures in the lateral temporal cortex of a different study cohort[15]. However, in work by others, no such relationship has been observed in the entorhinal cortex [64][67]. Such a contrast in relationships may relate to the regions selected for investigation. The entorhinal cortex is one of the earliest affected cortical regions with tau pathology in AD, and thus presumably represents a cortical area of increased vulnerability to tau pathology. Such vulnerability may mean that tau pathology may be generated in an Aβ independent manner within the region as is the case of primary age-related tauopathy [12]. Nevertheless, within the prefrontal cortex, a region which does not demonstrate robust age related NFT tau pathology and is not burdened with NFTs until late into the Braak NFT staging criteria (Braak V-VI), such modest pre-tangle tau-pathology generated in this region may be largely dependent on Aβ mediated mechanism. Such a mechanism may become modified under pathological conditions, either upon reaching a critical threshold of intracellular Aβ accumulation and/or via seed component invasion.

Interestingly within the grey matter, although a positive relationship between intracellular Aβ and phospho-tau was observed in non-AD control cases, an inverse relationship was observed in AD cases. Whilst intracellular Aβ levels remained elevated compared to controls, there has been reports of a reduction of intracellular Aβ levels in line with the deposition of Aβ plaques in mice models [68] as well as in serial observations in Down’s syndrome brains [69] and cases of late stage NFT mediated neurodegeneration [67]. Such contradictions may be due to differences in the specific regions of investigations. However, it is equally plausible that elevation of intracellular Aβ precedes and indeed acts as a source for extracellular plaque deposition, with the excessive deposition of plaques at later stages subsequently reducing intracellular Aβ levels as observed in animal model [68]. In turn tau pathology may continue to grow due to the influence of seed component species.

In any case, excessive intracellular Aβ accumulation, is unlikely to be benign. The familial Osaka E639Δ APP mutant which produces non-fibril E22Δ Aβ gives rise to an accumulation of intracellular Aβ oligomers in the absence of plaques. In AD patients or mouse models carrying the Osaka mutation, pronounced cognitive impairments, cellular stress, synaptic spine loss and critically pathological tau phosphorylation and conformational changes are observed [22, 45]. Whilst several of the downstream cellular dysfunction may be independent of tau [70], these studies nevertheless highlight the induction of tau pathology via Aβ independent of plaque formation, in support of the observed correlation of intracellular Aβ and phospho-tau observed here.

Given emerging evidence from clinical Aβ antibody trials[71], which supporting the targeting of soluble fibrillar Aβ species to consequently reduce tau pathology, further understanding the degree of interaction between Aβ and tau will provide greater insight into the mechanisms of AD related pathogenesis. Equally, in light of the facilitation of fibril seeding by the existence of pre-existing tau phosphorylation/pathology in mice [72–74], the targeting of pre-tangle soluble tau elevations in late stage affected brain regions, may protect against tau seed infiltration as part of AD disease progression, and may provide an effective stalling of the condition.

## Conclusions

Collectively this study demonstrates the robust correlation of AT-8 reactive tau and Aβ in the frontal cortex of both non-AD controls and AD cases when measured biochemically. Given that such linear increases in Aβ plaques and AT-8 pathology is not observed when quantified via IHC, the study demonstrates the potential influence of non-plaque Aβ in the intra-regional generation of tau pathology. Specifically, the occurrence and accumulation of intracellular Aβ, in line with AD pathological progression, at least in part correlated with AT-8 pathology and thus may contribute to production of tau pathology. This finding is supportive of the amyloid cascade hypothesis, yet in late-stage AD cases such a relationship may be diminished, with additional factors contributing to tau pathology, at least within the frontal cortex. Critically, the observation of a localised relationship between Aβ and phospho-tau in cases with low Braak NFTs stages implies that there is a degree of regionally generated AD related pathology, which may be tolerated within a physiological range. Following the age-related accumulation of pathology, this regionally produced burden may prime the region for the invasion of seed competent forms of tau originating from connected regions, raising the burden beyond a critical threshold, and thus removing the necessity of Aβ for the propagation of tau pathology.

## Acknowledgements

issue for this study was provided by the Newcastle Brain Tissue Resource which is funded in part by a grant from the UK Medical Research Council (G0400074), by NIHR Newcastle Biomedical Research Centre awarded to the Newcastle upon Tyne NHS Foundation Trust and Newcastle University, and as part of the Brains for Dementia Research Programme jointly funded by Alzheimer’s Research UK and Alzheimer’s Society

