## Supplemental Table 1 for "Post-mortem AT-8 reactive tau species correlate with non-plaque Aβ levels in the frontal cortex of non-AD and AD brains"

| Case no | Sex | Age | PMD | Braak Stage | Thal | CERAD | NIAAA | McKeith | LBBraak | Use |
| --- | --- | --- | --- | --- | --- | --- | --- | --- | --- | --- |
| Control cases | | | | | | | | | | |
| 1 | M | 55 | 50 | 0 | 0 | 0 | Not AD | 0 | 0 | Biochem+TMA |
| 2 | M | 47 | 29 | 0 | 0 | 0 | Not AD | 0 | 0 | Biochem |
| 3 | F | 78 | 34 | 0 | 1 | 0 | Possible AD | 0 | 0 | Biochem+TMA |
| 4 | M | 60 | 60 | 0 | 1 | 0 | Possible AD | 0 | 0 | Biochem+TMA |
| 5 | M | 66 | 56 | 0 | 2 | 0 | Possible AD | 0 | 0 | Biochem+TMA |
| 6 | M | 65 | 55 | 0 | 2 | 0 | Possible AD | 0 | 0 | Biochem+TMA |
| 7 | F | 70 | 72 | 0 | 1 | 0 | Not AD | 2 | 0 | TMA+IHC |
| 8 | M | 72 | 39 | I | 1 | 0 | Possible AD | 0 | 0 | Biochem+TMA+IHC |
| 9 | F | 71 | 43 | I | 2 | 0 | Possible AD | 0 | 0 | Biochem+TMA |
| 10 | F | 92 | 39 | I | 2 | 0 | Possible AD | 0 | 0 | Biochem+TMA |
| 11 | M | 59 | 78 | I | 3 | 0 | Possible AD | 0 | 0 | Biochem+TMA |
| 12 | F | 81 | 19 | I | 3 | 0 | Possible AD | 0 | 0 | Biochem+TMA |
| 13 | M | 63 | 59 | I | 3 | 0 | Possible AD | 0 | 0 | Biochem+TMA |
| 14 | M | 85 | 85 | I | 2 | 0 | Possible AD | 0 | 0 | TMA+IHC |
| 15 | M | 81 | 43 | II | 0 | 0 | Not AD | 0 | 0 | Biochem+TMA+IHC |
| 16 | M | 80 | 16 | II | 0 | 0 | Not AD | 1 | 3 | Biochem+TMA+IHC |
| 17 | F | 94 | 82 | II | 1 | 0 | Possible AD | 0 | 0 | Biochem+TMA+IHC |
| 18 | M | 88 | 28 | II | 1 | 0 | Possible AD | 0 | 0 | Biochem+TMA |
| 19 | M | 97 | 23 | II | 2 | 0 | Possible AD | 0 | 0 | Biochem+TMA |
| 20 | M | 77 | 83 | II | 3 | 0 | Possible AD | 0 | 0 | Biochem+TMA |
| 21 | F | 96 | 24 | II | 3 | 0 | Possible AD | 0 | 0 | Biochem+TMA |
| 22 | M | 91 | 29 | II | 3 | 0 | Possible AD | 0 | 0 | Biochem+TMA |
| 23 | F | 97 | 21 | II | 2 | 1 | Possible AD | 0 | 0 | Biochem+TMA+IHC |
| 24 | F | 97 | 21 | II | 2 | 1 | Possible AD | 0 | 0 | TMA+IHC |
| 25 | F | 80 | 22 | II | 1 | 0 | Possible AD | 0 | 0 | IHC |
| 26 | F | 96 | 95 | III | 0 | 0 | Not AD | 0 | 0 | Biochem+TMA+IHC |
| 27 | F | 88 | 22 | III | 0 | 0 | Not AD | 0 | 0 | Biochem+TMA+IHC |
| 28 | F | 95 | 36 | III | 0 | 0 | Not AD | 0 | 0 | Biochem+TMA |
| 29 | F | 75 | 53 | III | 1 | 0 | Possible AD | 1 | 1 | Biochem+TMA |
| 30 | F | 89 | 49 | III | 1 | 0 | Possible AD | 1 | 3 | Biochem+TMA+IHC |
| 31 | M | 92 | 50 | III | 1 | 0 | Possible AD | 0 | 0 | Biochem+TMA |
| 32 | F | 89 | 34 | III | 2 | 0 | Possible AD | 0 | 0 | Biochem+TMA+IHC |
| 33 | F | 80 | 31 | III | 2 | 0 | Possible AD | 0 | 0 | Biochem+TMA+IHC |
| 34 | M | 81 | 82 | III | 2 | 1 | Possible AD | 0 | 0 | Biochem+TMA |
| 35 | M | 75 | 82 | IV | 3 | 2 | Probable AD | 0 | 0 | Biochem+TMA |
| 36 | F | 87 | 21 | IV | 3 | 2 | Probable AD | 0 | 0 | Biochem+TMA |
| 37 | M | 86 | 50 | IV | 4 | 2 | Probable AD | 0 | 0 | Biochem+TMA |
| 38 | M | 88 | 18 | IV | 4 | 2 | Probable AD | 0 | 0 | Biochem+TMA |
| 39 | M | 88 | 69 | IV | 4 | 2 | Probable AD | 0 | 0 | Biochem+TMA |
| AD cases | | | | | | | | | | |
| 40 | M | 74 | 26 | V | 4 | 3 | Definate AD | 0 | 0 | Biochem+TMA |
| 41 | F | 81 | 73 | V | 5 | 3 | Definate AD | 0 | 0 | Biochem+TMA |
| 42 | F | 86 | 5 | V | 5 | 3 | Definate AD | 0 | 0 | Biochem+TMA |
| 43 | M | 96 | 74 | V | 5 | 3 | Definate AD | 0 | 0 | Biochem |
| 44 | F | 93 | 64 | V | 5 | 3 | Definate AD | 0 | 0 | Biochem+TMA |
| 45 | F | 93 | 64 | V | 5 | 3 | Definate AD | 0 | 0 | TMA+IHC |
| 46 | F | 79 | 65 | VI | 4 | 3 | Definate AD | 1 | 3 | Biochem+TMA |
| 47 | F | 86 | 69 | VI | 5 | 3 | Definate AD | 0 | 0 | Biochem+TMA |
| 48 | M | 78 | 37 | VI | 5 | 3 | Definate AD | 4 | 0 | Biochem+TMA |
| 49 | F | 87 | 54 | VI | 5 | 3 | Definate AD | 0 | 0 | Biochem+TMA |
| 50 | M | 90 | 69 | VI | 5 | 3 | Definate AD | 0 | 0 | Biochem+TMA |
| 51 | F | 89 | 85 | VI | 5 | 3 | Definate AD | 1 | 2 | Biochem+TMA+IHC |
| 52 | F | 90 | 90 | VI | 5 | 3 | Definate AD | 0 | 0 | Biochem+TMA+IHC |
| 53 | M | 80 | 24 | VI | 5 | 3 | Definate AD | 0 | 0 | Biochem+TMA |
| 54 | F | 92 | 74 | VI | 5 | 3 | Definate AD | 0 | 0 | Biochem+TMA |
| 55 | M | 91 | 72 | VI | 5 | 3 | Definate AD | 0 | 0 | Biochem+TMA |
| 56 | F | 78 | 40 | VI | 5 | 3 | Definate AD | 1 | 2 | Biochem+TMA+IHC |
| 57 | F | 83 | 52 | VI | 5 | 3 | Definate AD | 0 | 0 | Biochem+TMA+IHC |
| 58 | F | 84 | 46 | VI | 5 | 3 | Definate AD | 0 | 0 | TMA+IHC |
| 59 | M | 85 | 29 | VI | 5 | 3 | Definate AD | 0 | 0 | TMA+IHC |
| 60 | F | 81 | 50 | VI | 5 | 3 | Definate AD | 0 | 0 | TMA+IHC |

**Supplemental table 1 – Individual case list for tissue used.** Each case was assigned an arbitrary case number (Case no) and sex, age (years), post-mortem delay (PMD, hrs), Braak stage, Thal phase, Consortium to Establish a Registry for Alzheimer’s Disease (CERAD), the National Institute of Ageing – Alzheimer’s Association (NIA-AA) criteria, Lewy body (LB) Braak stage and McKeith criteria and tissue use of either biochemical( Biochem) analysis, tissue microarray for plaque and AT-8 pathology (TMA) and immunohistochemistry for intracellular Aβ and AT-8 (IHC) is provided.
