## Supplemental Table 3 for "Post-mortem AT-8 reactive tau species correlate with non-plaque Aβ levels in the frontal cortex of non-AD and AD brains"

|  | Braak | Thal | CERAD | NIA-AA | Age | PMD | AT8 | Plaques |
| --- | --- | --- | --- | --- | --- | --- | --- | --- |
| Total cohort | | | | | | | | |
| AT8 | **;r=0.55 | **;r=0.78 | **;r=0.86 | **;r=0.86 | N.S | N.S |  | **;r=0.68 |
| Plaques | **;r=0.67 | **;r=0.82 | **;r=0.7 | **;r=0.78 | N.S | N.S | **;r=0.68 |  |
| Controls | | | | | | | | |
| AT8 | N.S | *;r=0.41 | **;r=0.68 | **;r=0.48 | *;r=0.36 | N.S |  | N.S |
| Plaques | N.S | **;r=0.67 | N.S | **;r=0.55 | N.S | N.S | N.S |  |
| Alzheimer's Disease | | | | | | | | |
| AT8 | N.S | N.S | N.A | N.A | N.S | N.S |  | N.S |
| Plaques | N.S | N.S | N.A | N.A | N.S | N.S | N.S |  |

**Supplemental table 3. Correlation matrix of immunohistochemical measures of Aβ plaques and AT-8 reactive tau with neuropathological assessment.** Spearman correlations (r) between AT-8 and Aβ and Braak NFT staging, Thal phase, CERAD, NIA-AA, Age and postmortem delay (PMD) are shown. Note for Braak stage correlations reported are controlled for the influence of age. N.S = not significant, *=p<0.05, and **=p<0.01.
