## Supplemental Table 4 for "Post-mortem AT-8 reactive tau species correlate with non-plaque Aβ levels in the frontal cortex of non-AD and AD brains"

|  | Biochemistry | | | | TMA | | IHC | | | |
| --- | --- | --- | --- | --- | --- | --- | --- | --- | --- | --- |
|  | AT8-GM | AT8-WM | Total-Aβ-GM | Total-Aβ-WM | AT8 | Plaques | AT8-GM | AT8-WM | Intracell-Aβ-GM | Intracell-Aβ-WM |
| Total cohort | | | | | | | | | | |
| AT8-GM |  | **;r=0.71 | **;r=0.78 | **;r=0.57 | **;r=0.74 | **;r=0.64 | **;r=0.75 | N.S | **;r=0.74 | *;r=0.53 |
| AT8-WM | **;r=0.71 |  | **;r=0.54 | **;r=0.75 | **;r=0.68 | **;r=0.45 | **;r=0.73 | **;r=0.63 | **;r=0.75 | *;r=0.56 |
| Total-Aβ-GM | **;r=0.78 | **;r=0.54 |  | **;r=0.61 | **;r=0.67 | **;r=0.79 | **;r=0.79 | N.S | **;r=0.7 | *;r=0.63 |
| Total-Aβ-WM | **;r=0.57 | **;r=0.75 | **;r=0.61 |  | **;r=0.53 | **;r=0.59 | **;r=0.73 | **;r=0.69 | **;r=0.7 | *;r=0.57 |
| AT8 | **;r=0.74 | **;r=0.68 | **;r=0.67 | **;r=0.53 |  | **;r=0.68 | **;r=0.72 | **;r=0.73 | **;r=0.61 | **;r=0.59 |
| Plaques | **;r=0.64 | **;r=0.45 | **;r=0.79 | **;r=0.59 | **;r=0.68 |  | **;r=0.67 | *;r=0.46 | **;r=0.58 | N.S |
| AT8-GM | **;r=0.75 | **;r=0.73 | **;r=0.79 | **;r=0.73 | **;r=0.72 | **;r=0.67 |  | **;r=0.83 | **;r=0.78 | **;r=0.54 |
| AT8-WM | N.S | **;r=0.63 | N.S | **;r=0.69 | **;r=0.73 | *;r=0.46 | **;r=0.83 |  | **;r=0.74 | **;r=74 |
| Intracell-Aβ-GM | **;r=0.74 | **;r=0.75 | **;r=0.7 | **;r=0.7 | **;r=0.61 | **;r=0.58 | **;r=0.78 | **;r=0.74 |  | **;r=0.74 |
| Intracell-Aβ-WM | *;r=0.53 | *;r=0.56 | *;r=0.63 | *;r=0.57 | **;r=0.59 | N.S | **;r=0.54 | **;r=74 | **;r=0.74 |  |
| Controls | | | | | | | | | | |
| AT8-GM |  | **;r=0.48 | **;r=0.61 | N.S | *;r=0.4 | *;r=0.38 | N.S | N.S | N.S | N.S |
| AT8-WM | **;r=0.48 |  | *;r=0.37 | **;r=0.66 | **;r=0.52 | N.S | N.S | N.S | N.S | N.S |
| Total-Aβ-GM | **;r=0.61 | *;r=0.37 |  | **;r=0.59 | *;r=0.4 | **;r=0.61 | N.S | N.S | N.S | N.S |
| Total-Aβ-WM | N.S | **;r=0.66 | **;r=0.59 |  | N.S | *;r=0.49 | N.S | N.S | N.S | N.S |
| AT8 | *;r=0.4 | **;r=0.52 | *;r=0.4 | N.S |  | N.S | N.S | N.S | N.S | N.S |
| Plaques | *;r=0.38 | N.S | **;r=0.61 | *;r=0.49 | N.S |  | N.S | N.S | N.S | N.S |
| AT8-GM | N.S | N.S | N.S | N.S | N.S | N.S |  | **;r=0.72 | **;r=0.82 | N.S |
| AT8-WM | N.S | N.S | N.S | N.S | N.S | N.S | **;r=0.72 |  | **;r=0.71 | **;r=0.76 |
| Intracell-Aβ-GM | N.S | N.S | N.S | N.S | N.S | N.S | **;r=0.82 | **;r=0.71 |  | *;r=0.63 |
| Intracell-Aβ-WM | N.S | N.S | N.S | N.S | N.S | N.S | N.S | **;r=0.76 | *;r=0.63 |  |
| Alzheimer's Disease | | | | | | | | | | |
| AT8-GM |  | **;r=0.71 | **;r=0.78 | **;r=0.56 | *;r=0.57 | N.S | N.S | N.S | N.S | N.S |
| AT8-WM | **;r=0.71 |  | **;r=0.54 | **;r=0.75 | N.S | N.S | N.S | N.S | N.S | N.S |
| Total-Aβ-GM | **;r=0.78 | **;r=0.54 |  | **;r=0.6 | N.S | N.S | N.S | N.s | N.S | N.S |
| Total-Aβ-WM | **;r=0.56 | **;r=0.75 | **;r=0.6 |  | N.S | N.S | N.S | N.S | N.S | N.S |
| AT8 | *;r=0.57 | N.S | N.S | N.S |  | N.S | N.S | *;r=0.75 | N.S | N.S |
| Plaques | N.S | N.S | N.S | N.S | N.S |  | N.S | N.S | N.S | N.S |
| AT8-GM | N.S | N.S | N.S | N.S | N.S | N.S |  | N.S | N.S | N.S |
| AT8-WM | N.S | N.S | N.S | N.S | *;r=0.75 | N.S | N.S |  | N.S | N.S |
| Intracell-Aβ-GM | N.S | N.S | N.S | N.S | N.S | N.S | N.S | N.S |  | N.S |
| Intracell-Aβ-WM | N.S | N.S | N.S | N.S | N.S | N.S | N.S | N.S | N.S |  |

**Supplemental table 4. Correlation matrix of all measures of Aβ and AT-8 reactive tau across the study** Spearman correlations (r) between AT-8 and Aβ and Braak NFT staging, Thal phase, CERAD, NIA-AA, Age and post-mortem delay (PMD) are shown. Note for Braak stage correlations reported are controlled for the influence of age. N.S = not significant, *=p<0.05, and **=p<0.01.
