## Supplemental Table 5 for "Post-mortem AT-8 reactive tau species correlate with non-plaque Aβ levels in the frontal cortex of non-AD and AD brains"

|  | Braak | Thal | CERAD | NIA-AA | Age | PMD | AT8-GM | AT8-WM | Intracell-Aβ-GM | Intracell-Aβ-WM |
| --- | --- | --- | --- | --- | --- | --- | --- | --- | --- | --- |
| Total cohort | | | | | | | | | | |
| AT8-GM | **;r=0.62 | **;r=0.67 | **;r=0.76 | **;r=0.66 | N.S | N.S |  | **;r=0.83 | **;r=0.78 | **;r=0.54 |
| AT8-WM | **;r=0.5 | **;r=0.58 | **;r=0.7 | **;r=0.56 | N.S | N.S | **;r=0.83 |  | **;r=0.74 | **;r=74 |
| Intracell-Aβ-GM | **;r=0.61 | **;r=0.66 | **;r=0.75 | **;r=0.66 | N.S | N.S | **;r=0.78 | **;r=0.74 |  | **;r=0.74 |
| Intracell-Aβ-WM | N.S | *;r=0.5 | *;r=0.54 | *;r=0.42 | N.S | N.S | **;r=0.54 | **;r=0.74 | **;r=0.74 |  |
| Controls | | | | | | | | | | |
| AT8-GM | N.S | N.S | N.S | N.S | N.S | N.S |  | **;r=0.72 | **;r=0.82 | N.S |
| AT8-WM | N.S | N.S | N.S | N.S | N.S | N.S | **;r=0.72 |  | **;r=0.71 | **;r=0.76 |
| Intracell-Aβ-GM | N.S | N.S | N.S | N.S | N.S | N.S | **;r=0.82 | **;r=0.71 |  | *;r=0.63 |
| Intracell-Aβ-WM | N.S | N.S | N.S | N.S | N.S | N.S | N.S | **;r=0.76 | *;r=0.63 |  |
| Alzheimer's Disease | | | | | | | | | | |
| AT8-GM | N.S | N.A | N.A | N.A | N.S | N.S |  | N.S | N.S | N.S |
| AT8-WM | N.S | N.A | N.A | N.A | N.S | N.S | N.S |  | N.S | N.S |
| Intracell-Aβ-GM | N.S | N.A | N.A | N.A | N.S | N.S | N.S | N.S |  | N.S |
| Intracell-Aβ-WM | N.S | N.A | N.A | N.A | N.S | N.S | N.S | N.S | N.S |  |

**Supplemental table 5. Correlation matrix of immunohistochemical measures of intracellular Aβ and AT-8 reactive tau..** Spearman correlations (r) between AT-8 and Aβ and Braak NFT staging, Thal phase, CERAD, NIA-AA, Age and postmortem delay (PMD) are shown for both the grey matter (GM) and white matter (WM) samples. Note for Braak stage correlations reported are controlled for the influence of age. N.S = not significant, *=p<0.05, and **=p<0.01.
